# *In silico* approaches to identify the role of lncRNAs in Prostate cancer and Androgen receptor-targeted proteins

**DOI:** 10.1101/2023.10.07.561330

**Authors:** Barkha Khilwani, Bhumandeep Kour, Nidhi Shukla, Bhargavi R, Sugunakar Vure, Abdul S. Ansari, Nirmal K. Lohiya, Renuka Suravajhala, Prashanth Suravajhala

**Author notes:** Equal contributing authors (Barkha Khilwani, Bhumandeep Kour).

## Abstract

Long non-coding RNA (lncRNAs) are known to have a role in pathogenesis of a broad spectrum of malignancies.These are found to have a significant role as signal transduction mediators in cancer signaling pathways. Prostate Cancer (PCa) is emerging with increasing cases worldwide even as advanced approaches in clinical diagnosis and treatment of PCa are still challenging to address. To enhance patient stratification, there is an indefatigable need to understand risk that can allow new approaches of treatment based on prognosis. While PCa is known to have mediated androgen receptor (AR) stimulation, the latter plays a key role in regulating transcription of genes via nuclear translocation which in turn leads to response to androgens. LncRNAs have been implicated in developing clinical diagnostic and prognostic biomarkers in a broad spectrum of cancers. In our present study, 12 lncRNAs identified from clinical samples from our erstwhile PCa patients were docked with PCa and AR targeted 36 proteins. We identified three lncRNAs, *viz*. SCARNA10, NPBWR1, ANKRD20A9P are common between the targeted proteins and discern that SCARNA10 lncRNA could serve as a prognostic signature for PCa and AR biogenesis. We also sought to check the coding potential of interfacial residues associated with lncRNA docking sites.x

## Introduction

Prostate cancer (PCa) is the second most common malignancy among males and it is estimated that by 2040 there will be an approximate 1,017,712 new cases of PCa worldwide (Siegel et al., 2020, Gupta et al., 2020). With an increasing number of cases, multiple management options have been developed for diagnosis and treatment of PCa. These advances have enabled patient stratification by risk measurement and allow devising strategy of treatment based on prognosis and patient preferences (Rodrigues et al., 2012). PCa is dependent on androgen stimulation mediated by the androgen receptor (AR) even as AR plays a significant role in growth and differentiation of the healthy prostate (Koochekpour, 2010). For example, it is well known that the AR complexes move to the nucleus and dimerizes before modulating the transcription of targeted genes wherein many genes become regulatory and invite candidate genes to interact with. Furthermore, it also allows the drugs to be targeted on their AR grooves, thus making the active functional motifs.

The AR gene is located on the X-chromosome at the locus Xq11-12 and is composed of 8 exons coding for a ∼2757 bp open reading frame and ∼919 amino acids within a 10.6 kb mRNA (Brinkmann et al., 1989). The genomic structure of AR has been highly conserved throughout mammalian evolution. Similar to many other steroid receptors, the AR consists of distinct functional motifs organized as the amino-terminal domain (NTD; 555 amino acids coded by exon 1), DNA-binding domain (DBD; 68-amino acid coded by exon 2 and 3), ligand-binding domain (LBD; 295 amino acids coded by exons 4–8), nuclear localization (amino acid 628–657) and AF-1, AF-5 and AF-2 transactivation units encoded by exon 1 and 8, and a hinge region separates LBD from DBD. On the contrary, to verify high evolutionary conservation for LBD, DBD and the N-terminal of the hinge fragment, the most variable region is the NTD sequence. This domain is encoded by several regions of highly repetitive DNA sequences, such as CAG and GGC repeats. Similar to other nuclear receptors, the DBD region of AR contains nine cysteines, of which eight are linked to two zinc ions, and through the sulfhydryl groups they are organized in two zinc finger domains.Over the years a multitude of novel lncRNAs dysregulated in PCa had been identified by large scale RNA profiling projects which resulted in identifying a set of 121 PCa-associated intergenic non-coding RNA transcripts termed the PCAT family (Prensner et al., 2011). Functional analyses of lncRNAs have revealed significant contributions to PCa by targeting relevant pathways and gene regulation mechanisms including PTEN/AKT and AR signaling as well as chromatin remodeling complexes. AR is a member of the steroid and nuclear receptor superfamily of nuclear transcription factor (NR3C4, nuclear receptor subfamily 3, group C, gene 4) and is involved in the regulation of normal growth and development of various target organs (Gao et al. 2005). The AR is a ligand-dependent transcription factor, upon binding with androgens, i.e. native ligands 5α-dihydrotestosterone (DHT) and testosterone, AR initiates male sexual development and differentiation. In a nutshell, AR functions in response to androgens and regulates transcription of genes via nuclear translocation.

With the advancement of high throughput sequencing technologies such as next generation sequencing (NGS), our understanding of diseased phenotypes have increased, cancer being one of them. Our whole exome sequencing (WES) approach in PCa has identified ca. 30 causal genes in the Indian sub-population (Gupta et al., 2020). Similarly, the RNA-sequencing (RNA-seq) approach is used to measure the expression across a transcriptome and identify gene fusions, single nucleotide variants etc. In a parallel study in our lab, using the RNA-seq approach we have identified many differentially expressed genes (DEGs) and long non-coding RNAs (lncRNAs) (Shukla et al., 2023) in PCa samples specific to India. LncRNAs have been implicated as a diagnostic and prognostic biomarkers in different cancers (Qian et al., 2020) with well-known lncRNAs such as MALAT, HOTAIR, XIST have been reported in genitourinary cancers but their role as biomarkers is warranted (Tortora et al., 2023). From our study, we identified ca.12 lncRNAs, some of which are known to be involved in different cancers.

The lncRNAs play a very important role in the epigenetic regulation as nuclei primarily aid in gene transcriptional phase resulting in changes in DNA, histone, acetylation and methylation. We have earlier shown how lncRNAs play a very important role in the pathogenesis of various malignancies attributing to their role as key signal transduction mediators in cancer signaling pathways (Suravajhala et al., 2022). Association of lncRNAs with metabolic intermediates, proteins, signaling molecules, cellular lipids etc., facilitates intracellular signaling with an aid of lncRNAs and their target molecules correlating to tumor progression. Important role of lncRNAs has been identified in development of PCa, promotion of castration resistant PCa (CRPC), cell proliferation, invasion, metastatic spread along with modulation of AR-mediated signaling (Yang et al., 2021). On the other hand, AR signaling pathway involves various mediators regulated by lncRNA using various mechanisms, for example, Prostate cancer antigen 3 (PCA3) modulates PCa cell survival via modulating AR signaling and is now used in PCa diagnosis (Lemos et al., 2019), SChLAP1 (second chromosome locus associated with prostate-1) was identified as a highly prognostic lncRNA that differentially expressed in aggressive and indolent form of PCa (Presner et al., 2013). Considering the dynamic role of lncRNAs as novel prognostic, diagnostic and predictive markers in PCa, lncRNAs may also serve as therapeutic targets aiding in prevention, development and treatment of CRPC and metastasis of the disease. Functional analysis of lncRNAs could be done by deciphering lncRNA-protein interaction as the function of most lncRNAs is dependent on interaction with protein-coding genes. In contrast to limited experimental approaches available, lncRNA-Protein interactions can be more effectively studied employing different computational tools. It will be interesting to predict the lncRNA that are common in PCa and AR that can enable us to develop potential lncRNAs that are prognostic biomarkers. This study attempts to unfold the mechanisms involving lncRNA biogenesis and proteins associated with AR and PCa biogenesis.

## Materials and Methods

### PCa associated proteins, LncRNAs and Androgen Receptors (AR)

FASTA files of all the 28 PCa associated proteins, *viz. ADA, ANG, BRCA1, CTNS, HBB, GNPTAB, COL6A1, OTOF, TP53, CYP11B2, CYP1B1, GJB6, RHAG, DNAAF1, BRCA2, NF1 MCM8, MCCC1, CAPN3, MYO15A, MRE11, KRIT1, HEXB, SCN9A, PRLR, OPA1, ATP6V0A2 and USH2A* (Table 1) were retrieved from the NCBI (www.ncbi.nlm.nih.gov last accessed on May 25, 2023). The sequence data of of 11 lncRNAs (SCARNA10, LINC01973, LINC00940, NPBWR1, FLJ16779, ANKRD20A9P, LINC00298, SNHG19, LOC341056, TLX1NB, LINC00662:60) were extracted from NONCODE (http://www.noncode.org/ last accessed on May 25, 2023), LNCipedia (https://lncipedia.org/ last accessed on May 25, 2023 last accessed on May 25, 2023) and RNAcentral (https://rnacentral.org/) to get their HSAT ids’ and their respective FASTA files. The proteins were considered as receptors and LncRNAs as ligands with the FASTA files rendered as an input to the HDOCK server (Figure 1).

**Table 1.** List of PCa Proteins and PDB Ids used in the molecular docking study.

| PCa Proteins |  |  |
| --- | --- | --- |
| PROTEIN | PDB ID | REFERENCES |
| ADA | 3IAR | Ugochukwu et al., 2011 |
| ANG | 4AOH | Thiyagarajan et al., 2012 |
| ATP6V0A2 | 3RRK | Srinivasan et al., 2011 |
| **BRCA1** | 6GVW | Rabl et al., 2019 |
| **BRCA2** | 1MIU | Yang et al., 2002 |
| **CAPN3** | 6BGP | Ye et al., 2018 |
| **COL6A1** | 1KNT | Arnoux et al., 1995 |
| **CTNS** | 7ZKW | Löbel et al., 2022 |
| **CYP11B2** | 7M8I | Brixius-Anderko et al., 2011 |
| **CYP1B1** | 6OYV | Wang et al., 2021 |
| **GJB6** | 5ER7 | Bennett et al., 2016 |
| **GNPTAB** | 2N6D | Serrano et al., 2015 |
| **HBB** | 6LCX | Shibayama et al., 2020 |
| **HEXB** | 3LMY | Bateman et al., 2011 |
| **KRIT1** | 5D68 | Zhang et al., 2015 |
| **MCCC1** | 2EJM | Ruhul et al., 2007 |
| **MCM8** | 6L0O | Zeng et al., 2020 |
| **MRE11** | 3T1I | Park et al., 2011 |
| **MYO15A** | 7UDU | Gong et al., 2022 |
| **NF1** | 1NF1 | Naschberger et al., 2021 |
| **OPA1** | 6JTG | Yu C et al., 2020 |
| **OTOF** | 3L9B | Helfmann et al., 2011 |
| **PRLR** | 3NPZ | Van Angthoven et al., 2010 |
| **RHAG** | 8CSX | Vallese et al., 2022 |
| **SCN9A** | 7W9M | Huang et al., 2022 |
| **TP53** | 6VTC | Wu et al., 2020 |
| **Androgen Receptors** |  |  |
| **Androgen Receptor (AR)** | 4QL8 | Ullrich et al., 2014 |
| **Androgen receptor (AR)** | 2AM9 | Pereira de Jésus-Tran et al., 2006 |
| **Androgen receptor (AR)** | 2PNU | Cantin et al., 2007 |
| **Androgen receptor (AR)** | 1E3G | Matias et al., 2000 |
| **Selective Androgen Receptor Modulator (SARM)** | 5V8Q | Aikawa et al., 2017 |
| **Selective Androgen Receptor Modulator (SARM)** | 5CJ6 | Saeed et al., 2016 |
| **Uroporphyrinogen Decarboxylase (URO-D)** | 2Q71 | Phillips et al., 2007 |

**Figure 1.**
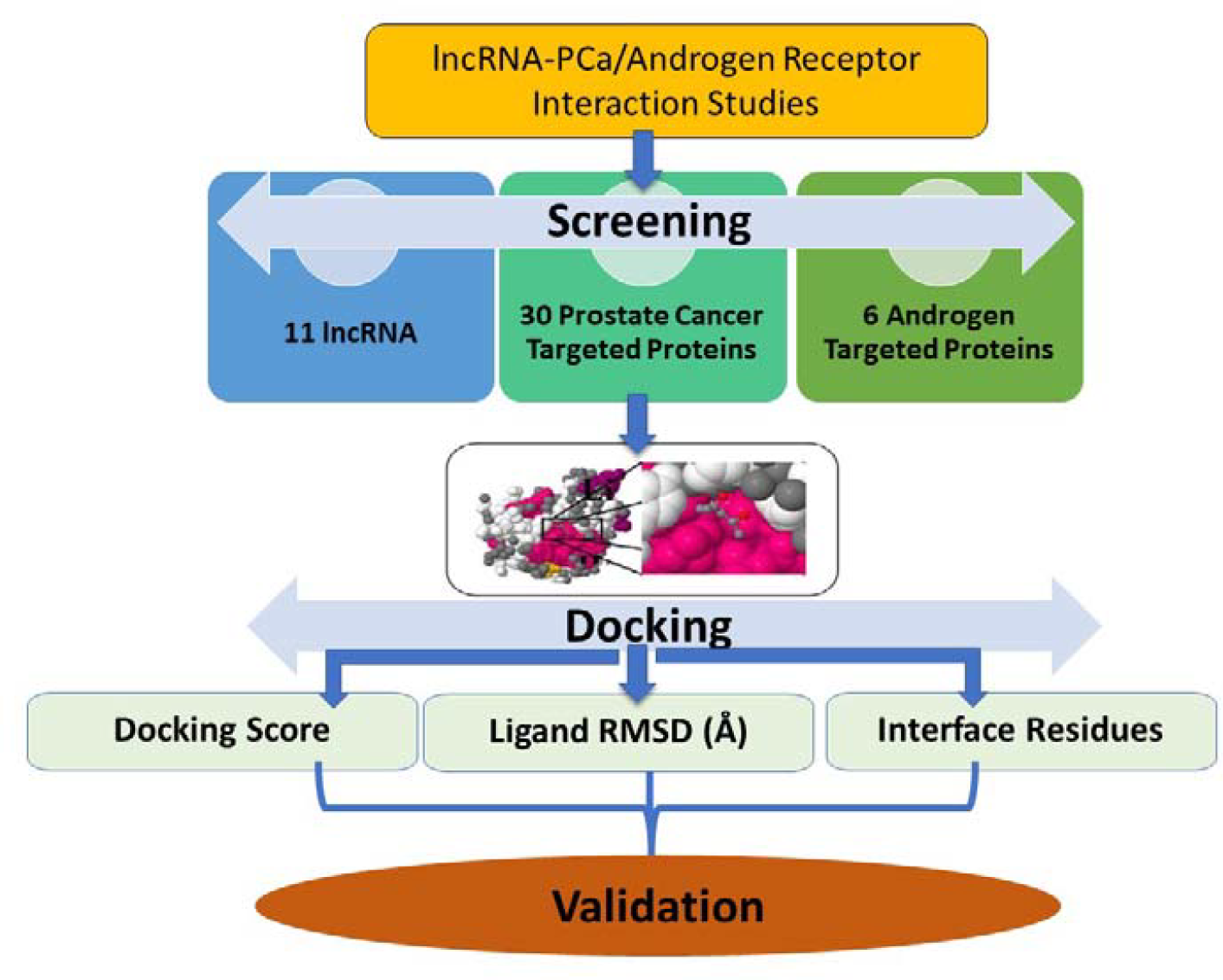
Systematic work flow of the 11 lncRNAs and the 36 targeted PCa and AR Proteins.

To further narrow down on lncRNAs, we checked whether or not any of these lncRNAs show any interaction with 30 causal genes obtained from our WES study. We used LncPro (http://cmbi.bjmu.edu.cn/lncpro last accessed on September 21, 2023) and RPI-Pred (http://ctsb.is.wfubmc.edu/projects/rpi-pred last accessed on September 21, 2023) for predicting the interaction between lncRNAs and protein-coding genes (supplementary files) and finally chose 5 lncRNAs which were showing strong interaction (above 90%) with some of the very important causal genes of PCa. For ARs (2Q71, 5V8Q, 4QL8, 2PNU, 5CJ6, 2AM9, 1E3G, 7KW7) we gave their protein data bank (PDB) id and chain ids as input. Both PDB and chain id were taken from RSCB PDB (https://www.rcsb.org/ t accessed on May 25, 2023). All the ARs were then considered as receptors and lncRNAs as ligands.

### Molecular docking studies

Docking analysis was performed using HDOCK (http://hdock.phys.hust.edu.cn/ last accessed on August 25, 2023), a docking tool. Numerous biological activities, including signal transmission, cell control, protein synthesis, DNA replication and repair, RNA transcription, etc., depend on interactions between nucleic acids and proteins. Thus, understanding their intricate structure will help researchers design treatment strategies or medications that specifically target these interactions. Because experimental approaches are expensive and technically challenging, molecular docking has become crucial in the identification of complex structures (Yan et al., 2017). For protein-protein docking, the HDOCK server (http://hdock.phys.hust.edu.cn/ last accessed on September 1, 2023) combines homology search, template-based modeling, structure prediction, macromolecular docking, biological information incorporation, and task administration. The server automatically predicts the interaction between receptor and ligand molecules using input data for both molecules (amino acid sequences or PDB structures). This was done using a hybrid method of template-based and template-free docking (Yan et al., 2020).

### Work flow of molecular docking

1. **Input:** Both protein sequences and structures were accepted as input data in the workflow’s initial step.The HDOCK server was built to take inputs for both protein sequences and structures, which makes it easier for both inexperienced and regular users to operate. The server takes two types of inputs for structures and two types of inputs for sequences for every molecule, given: 1) A PDB-formatted pdb file. ii) pdb file with chainID in PDB (e.g. 1CGI:E). iii) copy the protein sequence and paste it in the FASTA format. iv) Uploading a FASTA-formatted protein sequence file. Each molecule just requires one kind of input and with automated modeling of DNA/RNA structures from sequences currently difficult, the service only accepts structure inputs for DNAs and RNAs at this time (Yan et al., 2017).
2. **Sequencing similarity:** To determine the homologous sequences for both receptor and ligand molecules, this similarity match was carried out against the PDB sequence database using the sequences from input or converted from structures. The HHSuite software was used for protein sequence searches since it is widely known because of its effectiveness in locating distant homologs. Since FASTA (version 3.6) is a powerful and user-friendly tool for both protein and DNA/RNA sequence search, it is utilized for DNA/RNA and thus two sets of homologous templates are produced as a result of this process.
3. **Template selection:** After that, the process moves on to the third stage, which involves comparing two sets of templates to check if they shared any entries having the similar PDB codes. A similar template was chosen both for the receptor and the ligand if there are any such PDB codes. The best templates for the receptor protein and/or ligand protein were chosen from two sets of homologous templates, respectively, assuming that there was no link between the two sets. When there are many templates present, then the one with maximum sequence coverage, sequence similarity, and resolution is chosen. Models are constructed using MODELLER with the chosen templates, and ClustalW was used for sequence alignment.
4. **Result:** As the HDOCK server adds docking tasks to the queue on providing input three task status including “QUEUED,” “RUNNING,” and “RESULTS,” are yielded and finally the docking results found at http://hdock.phys.hust.edu.cn/date/jobid, where “jobid” is the specific job id displayed on the web page of status was retrieved.
5. **Output:** The docking output consists of three fundamental files: Receptor PDB file created by the server using the users’ FASTA sequence or supplied by users, Ligand PDB file created by the server using the user-provided FASTA sequence or supplied by users and ligand binding modes reflected by their transformations in the HDOCK output. Additionally, the result page displays a docking summary of the top 10 models at the bottom and the template information for the receptor and ligand at the top (Yan et al. 2017).

Based on the highest ligand receptor binding energy interacting model of each interaction was selected and these models were then subjected for visualization.

#### Visualization

The docked scores of lncRNA with proteins generated 180 complexes with 10 best poses for each lncRNA-protein resulting in 1800 models. Screening of the models was carried by least-bind energy complexes; 12 models of the complexes were considered to be identified of which, 5 complexes with PCa and 7 complexes with AR.The lowest energy known to have more stability were considered for 3D visualization with PyMOL software. The parameters were assigned to ligand site hydrogen bonds with <3 Bond distances were identified that indicate the high intensity and the possible orientation of the proteins resulting in the stable complex formation.

#### Coding potential

To check if there is any coding potential attributing to the interfacial residues, we performed checks using intrinsic feature estimation by employing CPC2 (Kang et al. 2017). The tool works on the premise that the ORF length coverage is estimated along with Fickett and hexamer scores which serve as a prudent classifier for estimating ncRNAs. The Fickett score is based on frequencies of A,T,G,Cs leaning upon the intrinsic divergence between ncRNAs.

## Results and Discussion

From our recently published RNA-Seq analyses (Shukla et al. 2023), 11 lncRNAs identified (SCARNA10, LINC01973, LINC00940, NPBWR1, FLJ16779, ANKRD20A9P, LINC00298, SNHG19, LOC341056, TLX1NB, LINC00662:60) were screened in the form of FASTA files which were then retrieved as discussed previously (using NCBI, NONCODE, LINCipedia and RNA central public portals). We considered the HDOCK web server for the molecular docking studies to identify the interactions of receptor-ligands. Among the 11 lncRNAs (Refer materials), 6 lncRNAs LINC01973, FLJ16779, LINC00298, SNHG19, LOC341056, LINC00662:60 with a limit of 5000 residues were not done as the interactions were not deciphered with no alternate tools available to consider. We performed docking on 5 lncRNAs (SCARNA10, LINC00940, NPBWR1, ANKRD20A9P and TLX1NB) with 28 PCa and AR targeted proteins. Five lncRNAs (SCARNA10, LINC00940, NPBWR1, ANKRD20A9P and TLX1NB) with 27 PCa associated proteins except USH2A resulted best confirmers each generated 10 models of which *CTNS (PDB ID: 5CTG);ANG(PDB ID: 4AOH);CYP1B1 (PDB ID:3PM0)* observed to show more stable complex formation (Table 2).

**Table 2:** The potential candidate 5-lncRNAs’ interactions with 3-PCa proteins (PDB id) and binding energies,root mean square deviation (RMSD) and Interacting residues.

| S.No. | LncRNA | PCa<br>Protein | PDB ID | Docking<br>Score | Ligand<br>RMSD<br>(Å) | Interface Residues |
| --- | --- | --- | --- | --- | --- | --- |
| 1 | TLX1NB | CTNS | 7ZKW | -322.82 | 57.44 | Ser177, Lys 176, Trp 304,<br>Asn 322, Asp 324, Phe 317,<br>Thr 261 |
| 2 | SCARNA10 | ANG | 4AOH | -259.38 | 482.89 | Gly72, Asn87, Lys89,<br>Ser99, Lys 97, Phy100 |
| 3 | NPBWR1 | ANG | 4AOH | -230.33 | 740.84 | Lys41, Gly72, Ser99,<br>Arg146 |
| 4 | ANKRD20A9<br>P | ANG | 4AOH | -220.15 | 803.24 | Asn 73, Ser 99, Lys 97 |
| 5 | LINC00940 | CYP1B<br>1 | 6OYB | -201.96 | 3095.39 | Gly72, Ser99, Ile143 |

### LncRNA-PCa interactions

The docked complexes of TLXINB-CTNS (PDB ID: 5CTG) with binding energy -322.82 kcal/mol and interacting/interfacial residues were identified, *viz*. Ser177,Lys 176, Trp 304,Asn 322,Asp 324, Phe 317,Thr 261 (Figure 2a); SCARNA10 -ANG(PDB ID: 4AOH) observed to have the least binding energy - 259.38 kcal/mol and interacting residues was identified as Gly72, Asn87, Lys89, Ser99, Lys 9, Phy100 (Figure 2b); in complex NPBWR1-ANG(PDB ID:4AOH) -230.33 kcal/mol and interacting residues was identified as Lys41, Gly72, Ser99, Arg146 (Figure 2c); in complex ANKRD20A9P-ANG (PDB ID: 4AOH)-220.15 kcal/mol and interacting residues was identified as Asn 73, Ser 99, Lys 97 (Figure 2d), while the LINC00940-CYP1B1 (PDB ID:3PM0) -201.96 kcal/mol and interacting residues were identified as Gly72, Ser99, Ile143 (Figure 2e) .

**Figure 2.**
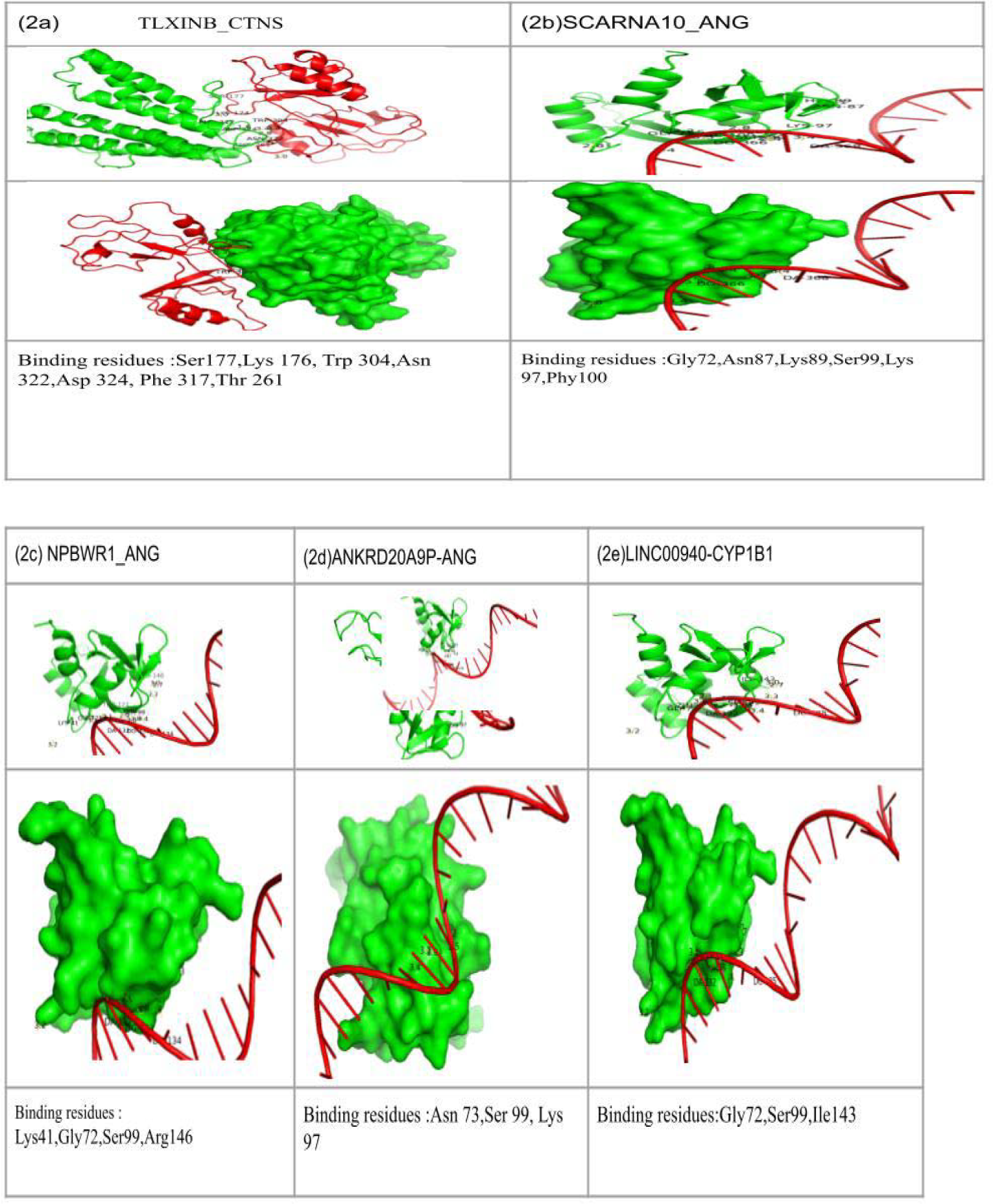
The 3D visualization using Pymol the complexes of potential 5-lncRNAs and interacting residues with 3-PCa proteins (2a-2e).

### LncRNA-AR interactions

SCARNA10, LINC00940, NPBWR1, ANKRD20A9P, TLX1NB were considered to know the possible interaction with AR targeted proteins (PDB id:2Q71, 5V8Q, 4QL8, 2PNU, 5CJ6, 2AM9, 1E3G, 7KW7) were considered. From the interaction studies,we identified least binding energy in 1E3G-ANKRD20A9P, 2AM9-SCARNA10, 5V8Q-NPBWR1 with -212--208 Kcal/mol binding energy as shown in Table 3. 1E3G-ANKRD20A9P has binding energy of -212.74 kcal/mol, with ligand RMSD of 742.99Å and residues are Arg-85,Trp-796,Arg-846 (figure 3a); 2AM9-SCARNA10 with binding energy of -208.93 kcal/mol, ligand RMSD as 536.64Å and the interaction residues are Thr755, Arg774, Leu-700 as shown in figure 3(3b); While 5V8Q-NPBWR1 with -208.04 kcal/mol, ligand RMSD 677.49Å of and interacting residues was identified as His-917(figure 3c); 2PNU-SCARNA10 with binding energy as -201.39 kcal/mol, RMSD 613.81Å and interacting residues Thr-850, Ser-853,Tyr-857(figure 3d); 5CJ6-SCARNA10 complex has binding energy of -197.78 kcal/mol, RMSD of 631.85Å and binding residues are Trp-796,Leu-797,Pro-868, Thr-918 (figure 3e; and 2Q71-SCARNA10 complex showed binding energy of -193.91 kcal/mol, RMSD of 634.84Å and its binding residues are Gln-919,Gln-792, Tyr-857(figure 3f) were visualized the 3D complexes with hydrogen interactions bellow 3 Å bond distance. While in 4QL8-ANKRD20A9P, we obtained the highest binding energy of -185.99kcal/mol with 762.89+ Å as ligand RMSD value. Our study indicates the SCARNA10 as a potential lncRNA having a lowest binding affinity energy of -259.38 with SCARNA10-ANG and -259.38 with SCARNA10-2AM9) suggesting their potency.

**Table 3:** The potential candidate 5-lncRNA interactions with 7-AR proteins (PDB id) and binding energies,RMSD and interacting residues.

| S.No. | LncRNA | Androgen Receptor Protein | Docking Score | Ligand RMSD (Å) | Interface Residues |
| --- | --- | --- | --- | --- | --- |
| 1 | ANKRD20A9P | 1E3G | -212.74 | 742.99 | Arg-85,Trp-796,Arg-846 |
| 2 | SCARNA10 | 2AM9 | -208.93 | 536.64 | Thr755, Arg774, Leu-700 |
| 3 | NPBWR1 | 5V8Q | -208.04 | 677.49 | His-917 |
| 4 | SCARNA10 | 2PNU | -201.39 | 613.81 | Thr-850, Ser-853,Tyr-857 |
| 5 | SCARNA10 | 5CJ6 | -197.78 | 631.85 | Trp-796,Leu-797,Pro-868, Thr-918 |
| 6 | SCARNA10 | 2Q71 | -193.91 | 634.84 | Gln-919,Gln-792, Tyr-857 |
| 7 | ANKRD20A9P | 4QL8 | -185.99 | 762.89 | Trp-796,Gln-792 |

**Figure 3.**
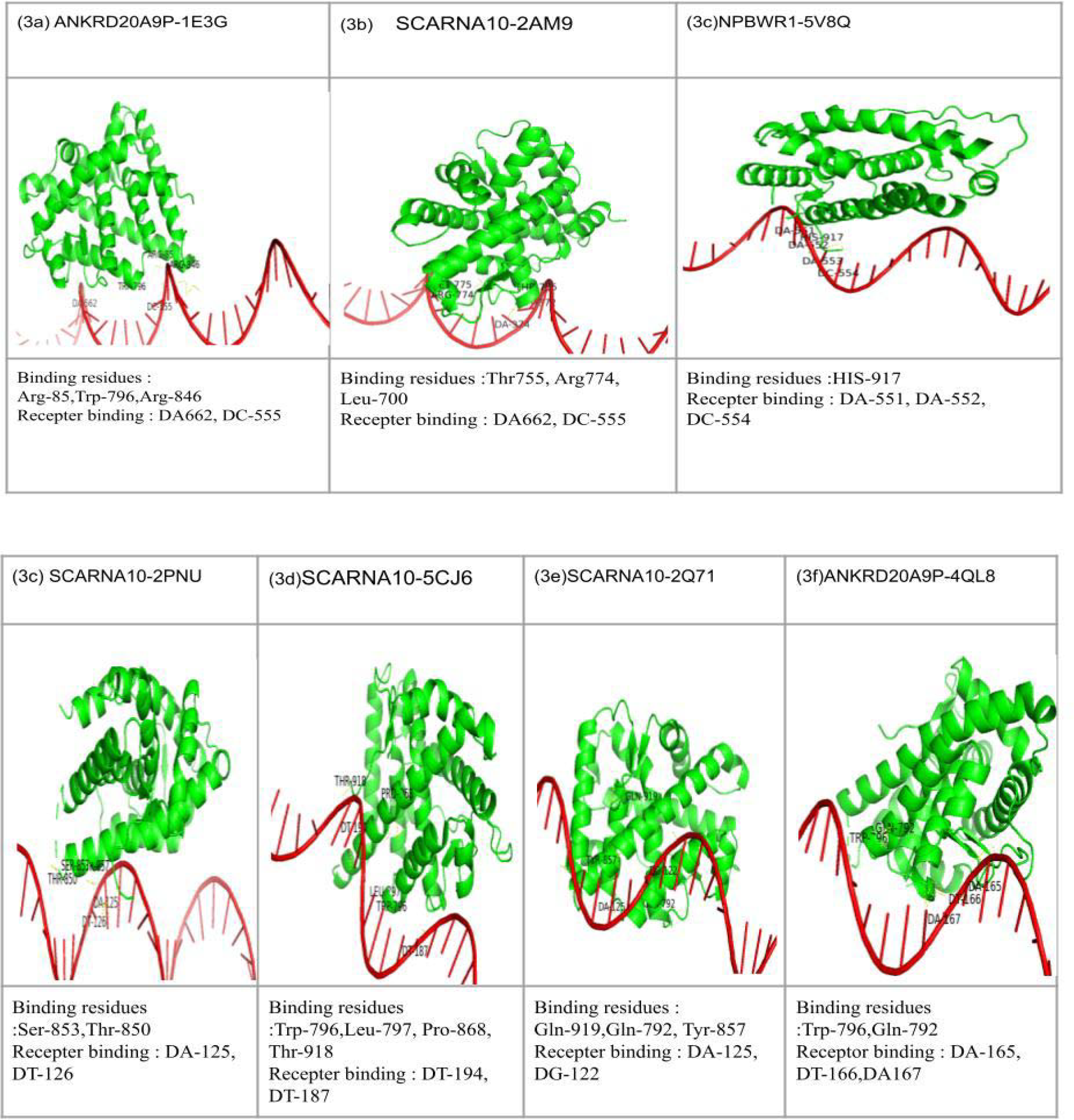
The 3D visualization using Pymol the complexes of potential 5-lncRNAs and interacting residues with 7 targeted AR proteins(3a-3f).

**Figure 4.**
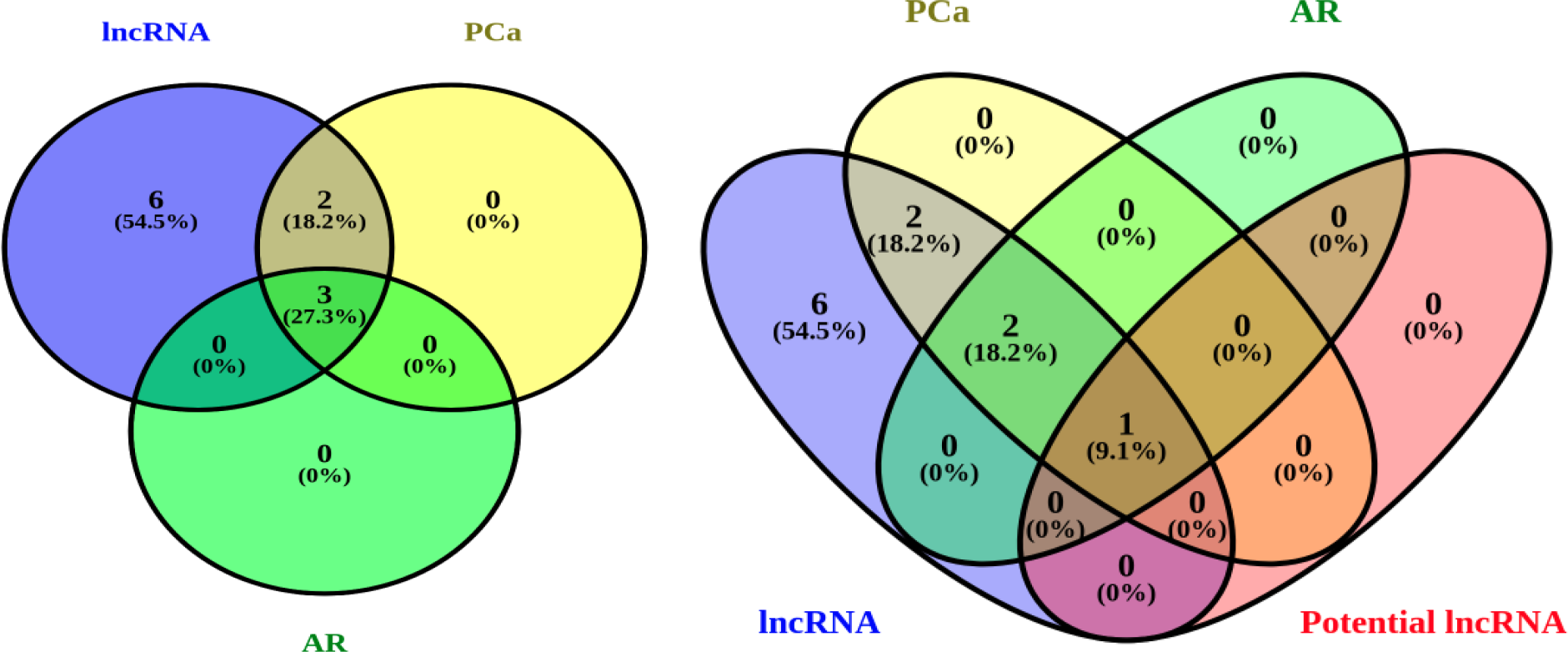
Statistical overview of 11 lncRNAs and 3 lncRNAs that are common in which SCARNA10 is considered as a potential biomarker .

**Figure 5:**
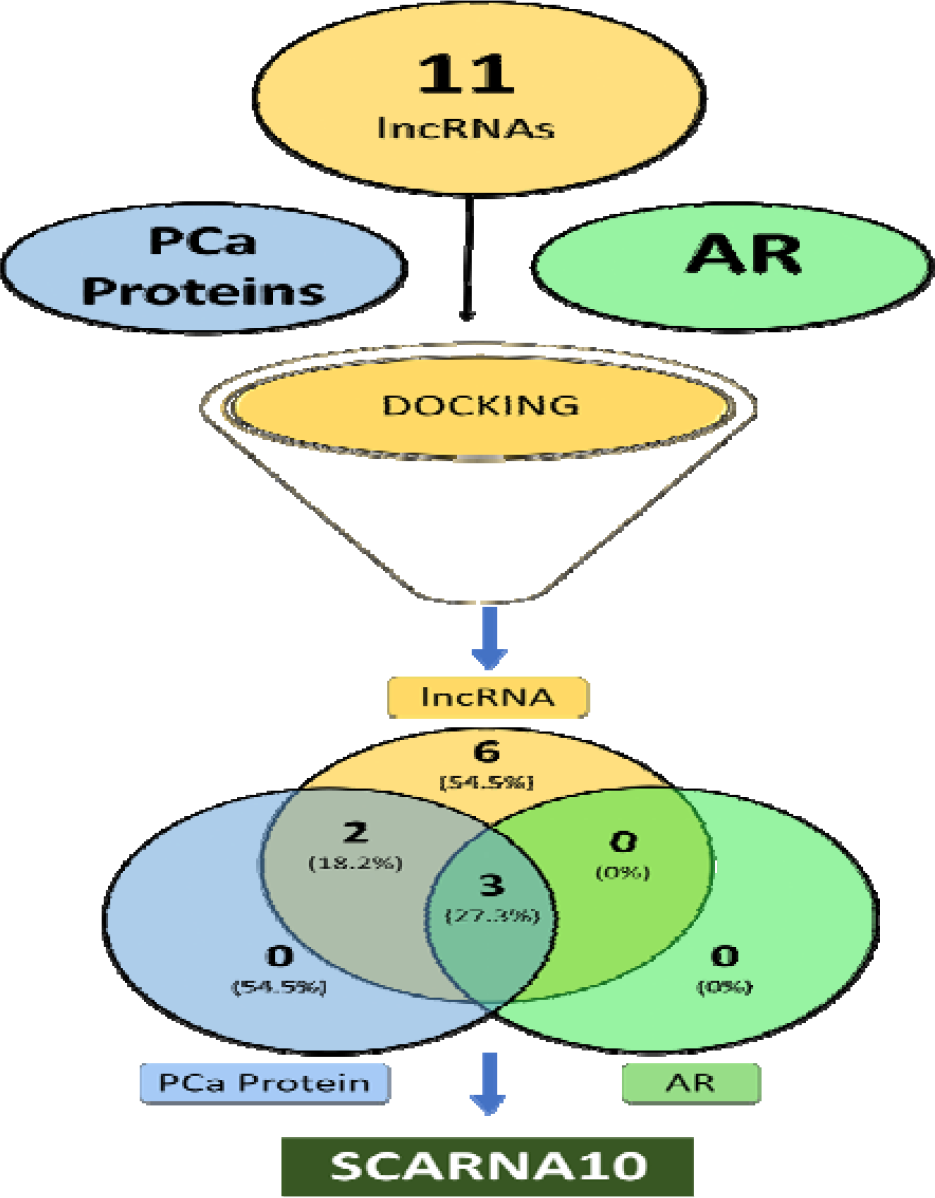
Summary of scoring results obtained after docking of 11 lncRNAs with PCa associated proteins and AR) following identification of SCARNA10 (lncRNA) as a common potential target.

**Figure 6:**
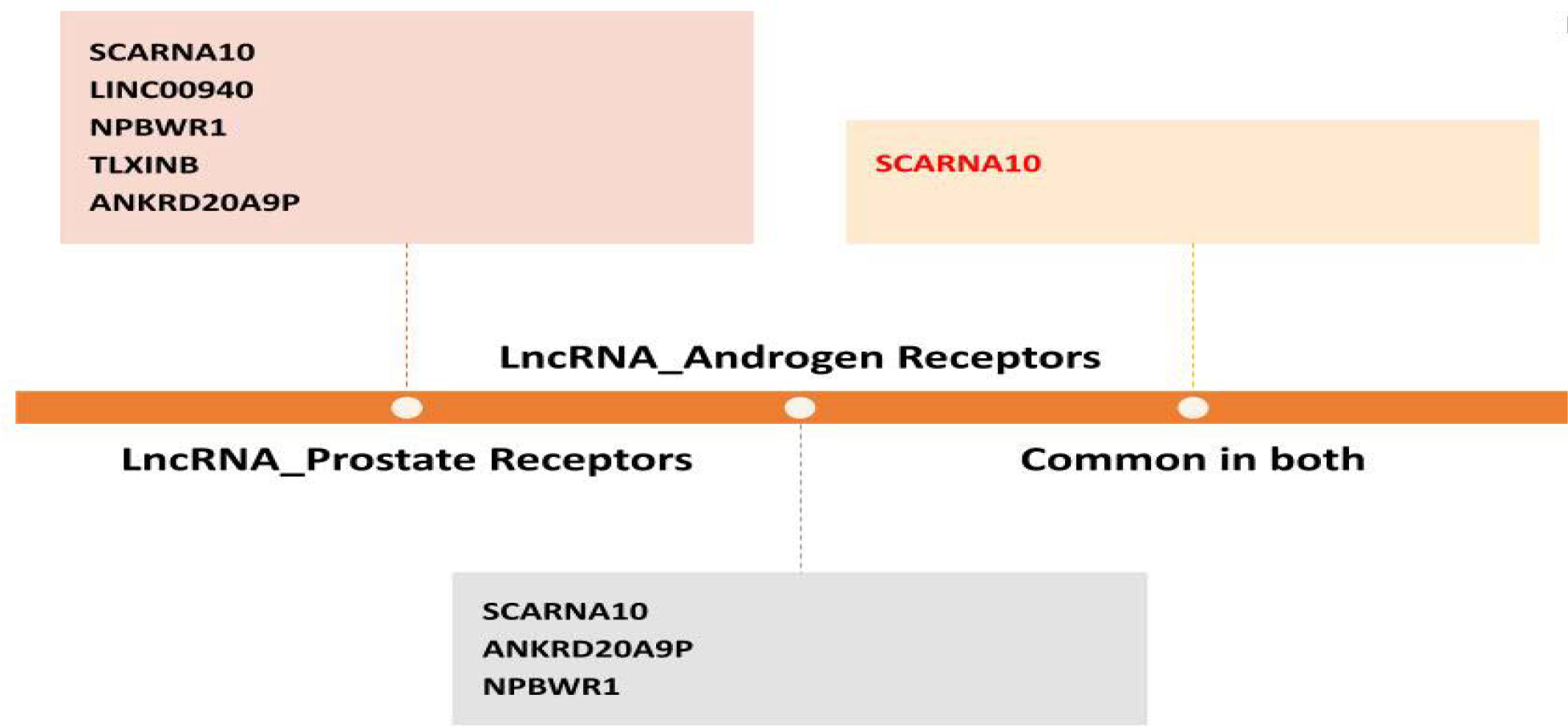
Overview of 11 lncRNAs interactions with 28 PCa proteins and 7 protein studies to identify potential lead SCARNA10 lncRNA that is common.

Recent studies have revealed that lncRNAs can regulate androgen signaling through various mechanisms (Kumar et al., 2021). It has been reported that lncRNAs can transactivate AR by binding to its enhancer region. LncRNAs are significant in prostate tumorigenesis as gene transcription regulatory sequences. PCGEM1, HOX transcript antisense RNA (HOTAIR), PCa gene 3 (PCA3) are some examples of lncRNAs that function as oncogenic and/or tumor suppressor in PCa through AR signaling pathway (Zang et al., 2016). Deficiency effect was reported in Androgen related transcription factor 1 (GDS5606 / 7953383) and carboxyl terminal-binding protein 2 in (FoxA1 Knockdown) studies, Peptidyl-prolyl cis/trans isomerase Pin1 (GDS5805 / 7953383) LNCaP. The phased mutation and SV Integrated studies reported that in LAPC4/LAPC4-CR cell lines duplications were found to increase due to CDK12 mutations (Wu et al., 2020). While PC3 and VCaP cells reported to have chromothripsis induced TP53 mutations, these reports indicate that high somatic mutations related to events in chromoplexy and chromothripsis have a critical role in PCa pathogenesis. Few limitations such as rearrangement were not considered in aneuploidy. In our lab, RNA-seq data from a small cohort of PCa patients was analyzed and some critical lncRNAs were identified as deterministic markers (Shukla et al., 2023). In present study, docking analysis of prostate specific proteins and AR with lncRNAs was performed to explore binding potential towards identifying interacting residues and putative RNA-binding motifs. Androgen signaling plays a vital role in PCa development and in treatment strategies (Jacob et al. 2021) as AR splice variants, amplification/overexpression of androgens, AR-Ligand binding stimulates development of CRPC. Our studies employed identifying target regions in AR to modulate and control AR-signaling pathways. Androgen dependent signaling instigates progression of PCa from benign to malignant disease with molecules like enzalutamide and abiraterone acetate playing a critical role as second-generation anti-androgen therapy (Efstathiou et al. 2020).

From the docking studies, SCARNA10, a candidate lncRNA common to both the PCa and AR targeted proteins was identified as While it is known to inhibit targeted gene binding of a promoter to polycomb repressive complex 2 (PRC2) suppressing the TGF signaling, we aimed to understand the mechanism of SCARNA10, when silenced, reducing the levels of TGF, TGF R1, KLF6 and Smad2,3 (Ganguly et al., 2021; Mao et al., 2020). It was also reported that SCARNA10 has a major role in chromosomal mutation at 12p13.31 identified in PCa metastasis studies (Saaid et al., 2014). In a few studies, SCARNA10 was reported to be up-regulated in breast and lung cancer (Han et al., 2022). Very limited data was reported of which the NCBI/GEO datasets reported that SCARNA 10 in PCa considering LNCaP cell line. Furthermore, it was identified that mR-135b, FOXA1 (GDS4957 / ILMN_3246209; LNCaP.FoxA1.6) were found to be overexpressed in clinical human PCa samples,VprBP depletion effect (GDS4829 / ILMN_3246209) was reported. The studies provide potential lncRNA prognostic biomarkers for future PCa research.

### Coding the uncoded

We observed that NONHSAT032215.2 is a human non-protein coding lnc-TUBA3C-16:1 gene and NONHSAT126578.2 have complete ORFs with putative peptides while NONHSAT026096.2 is with an incomplete peptide indicating even as the probability of coding potential is par 5%. We argue that these regions could be at the interfacial sites and serve as candidate regions for selective epitope binding (Table 4).

**Table 4:**
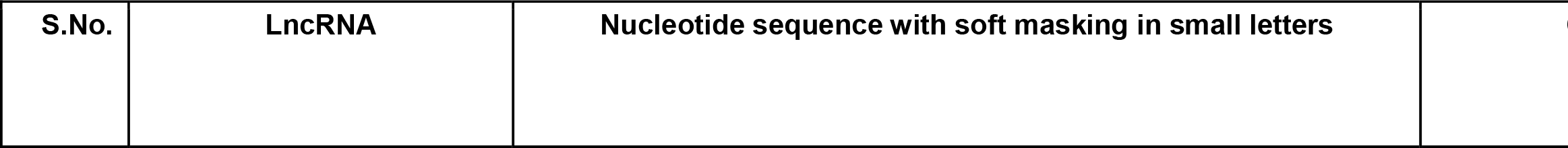
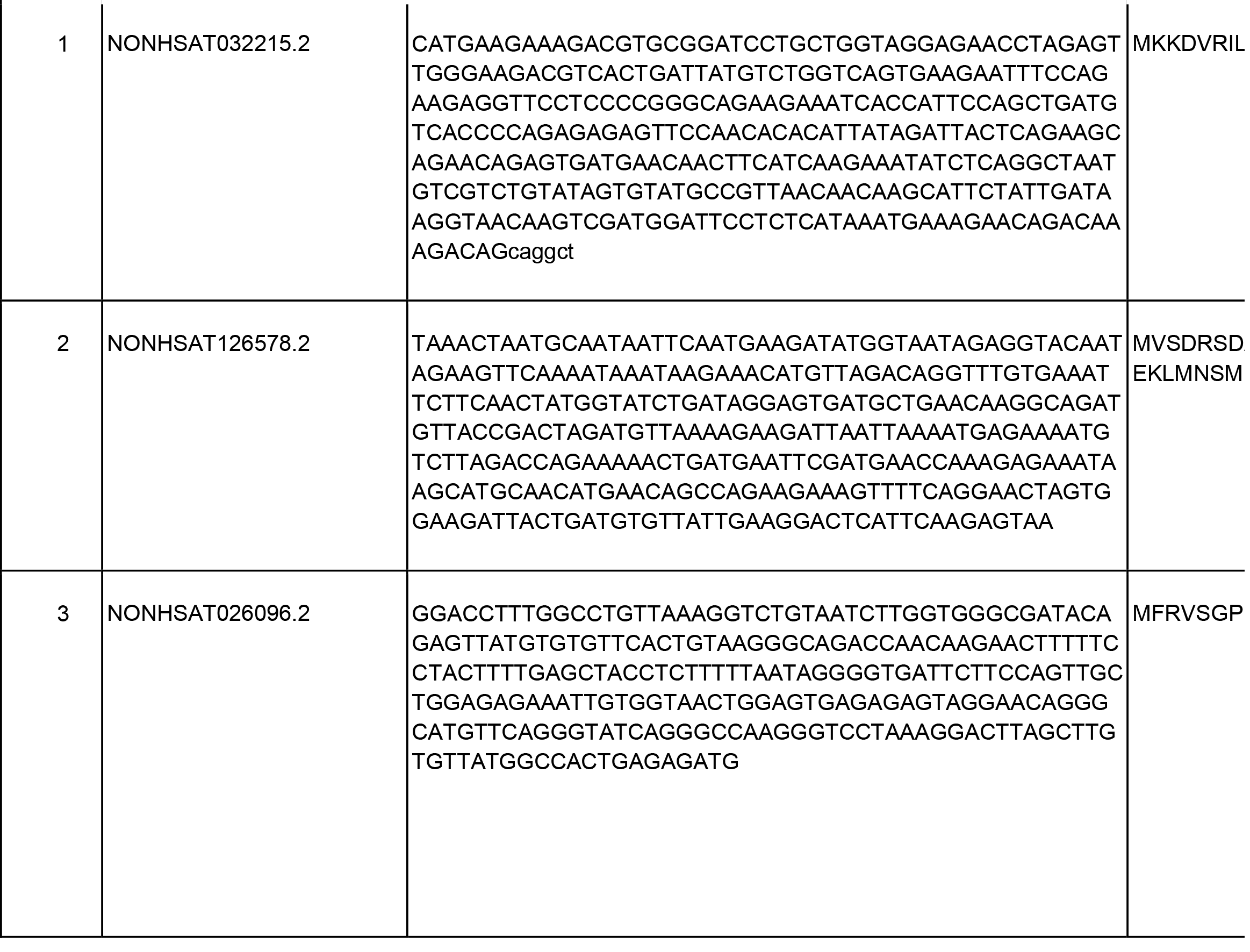
Coding potential of lncRNAs along with their putative peptides.

What remains intriguing is the coding potential of lncRNAs as we have considered those potential coding peptides and looked for similarity searches. For example, NONHSAT032215.2 was found to have similarity to mitochondrial Rho GTPase 1 isoform and RHOT genes and these potential peptides were indeed suppressed over the evolution. Deregulated Rho GTPases have been discovered in various tumors, including prostate which are known to promote the metastatic properties of human cancer cells (Jung et al. 2020). Similarly, Histone acetyltransferase 1 is known to upregulate AR expression to modulate CRPC cell resistance to well known drugs such as enzalutamide (Hong et al. 2021) and recently it is shown to epigenetically influence the AR signaling (Nguyen et al. 2023)

## Conclusions

With the advent of NGS approaches, identifying lncRNAs among the DEGs play a crucial role to bridge the gap between regulatory mechanisms of PCa and AR. In this work, we attempted to dock lncRNAs discovered from our previous PCa analysis to that of AR and infer candidate interactions. While we identified SCARNA10 as common lncRNA between both PCa and AR and argue that the functional aspect of this lncRNA would give us some insights into PCa progression, there is a room for identifying putative prognostic signatures for PCa detection. We also sought to ask whether or not any interfacial residues are at the site of coding potential of lncRNAs. As a future perspective, we envisage identifying specific lncRNAs that play crucial roles in AR signaling which could lead to the development of targeted therapies and further perform *in vitro* validation. Modulating the activity of these lncRNAs could potentially be a novel approach for treating AR-related diseases. Furthermore, understanding their interactions with AR could lead to the development of diagnostic or prognostic tools. If specific lncRNAs are found to be critically involved in AR signaling, we believe that this knowledge could be applied in the context of personalized medicine. Tailoring treatments based on a patient’s unique genetic and molecular profile could lead to more effective therapeutic strategies. Nonetheless, there could be microRNAs as well (miRs) at the helm of these interfacial sites, but as lncRNAs are largely regulatory, we deem to bridge the gap between lncRNAs and proteins associated with AR signaling.

## Authors’ contributions

BK, RS, NKL, AS, PS conceived the project. BK and BdK contributed equally to the project. NS, SV and BR identified the lncRNAs, PS ideated the project with NKL, supervised and curated the project, and proofread the manuscript before all the authors agreed to final submission.

## Acknowledgements

PS and RS gratefully acknowledge the chancellor, revered Mata Amritanandamayi Devi and Bipin G Nair, Dean, Amrita School of Biotechnology, Amrita Vishwa Vidyapeetham for their support. The authors thank the CAPCI consortium members on the project that they are associated with.

## Data availability

None

